# Rhythmic motor activity alleviates auditory attentional blinks

**DOI:** 10.64898/2026.05.05.722900

**Authors:** Zhiyuan Xu, Jiaqiu Vince Sun, Yuhan Lu, Wanting Zhang, Zhaoxin Wang, Yixuan Ku, Xing Tian

## Abstract

Effective sensory processing relies on both attention and the motor system, yet whether motor activity could provide attention-like functions to regulate perception remains unknown. We hypothesized that rhythmic motor signals could provide phasic regulation of prioritizing and sampling perceptual targets. Using an auditory attentional blink paradigm that created a temporal deficit in selective attention, we found that temporally aligned finger tapping improved the probe detection during the attentional blink window but impaired performance when attentional resources were abundant. Furthermore, transcranial alternating current stimulation (tACS) over the right sensorimotor cortex alleviated attentional blink when the probe was close to the peak of the stimulation, whereas stimulation over the left aggravated attentional blink when the probe was close to the trough. These results suggest that the motor system is a resource-dependent rhythmic regulator of attentional sampling. Motor signals can override attentional bottlenecks, suggesting the motor system as an active shaper of cognitive processes.

## INTRODUCTION

Our brain continuously receives sensory input from the external world as well as reafference from our own actions. The ability to effectively select and process sensory inputs across time is crucial to guide our behavior. Attention has been attributed to such optimization by selectively prioritizing neural representations that are most related to current behavioral goals (Buschman & Kastner, 2015). The motor system, classically viewed as the output end of cognition, also influences sensory processing. By sending internal signals, termed as efference copy and corollary discharge, to the sensory pathway, the motor system regulates the sensory system to update and predict the self-generated states during actions (Crapse & Sommer, 2008; S. Li et al., 2020; Rao, 2024; Tian & Poeppel, 2010; Wolpert et al., 1995). Given that both attention and the motor system actively constrain sensory processing, could they be attributed to a shared underlying mechanism? That is, can motor activity induce attention-like effects?

Rhythmicity is a fundamental property of the motor system and underpins various cognitive functions, especially those involving temporal processing, such as beat-based timing (Teki et al., 2011), time estimation (J. Coull & Nobre, 2008), and the internal clock (Strassmann et al., 2026) mechanisms. A primary function of attention is to orient (Posner, 1980, 2016) or shift (Buschman & Kastner, 2015; Fiebelkorn & Kastner, 2019) the ‘spotlight’ to the target. The motor system has been demonstrated for orienting attention across time – a function that uses temporal regularity to predict and select targets relevant to behavior goals (Morillon et al., 2015; Schroeder et al., 2010). For example, overt rhythmic movement sharpens auditory perception in the context of speech and music (Morillon et al., 2014; Zalta et al., 2020). The motor-rhythm-based effects of temporal orientation were observed in the left sensorimotor cortex encoded by oscillatory activity in the delta and beta bands, while the general benefits of movement are associated with the right sensorimotor cortex (Morillon & Baillet, 2017). These findings suggest a functional link between auditory attention and the motor system, implying that the sensorimotor circuits may provide a mechanistic pathway for temporally shifting attention.

The deterministic aspect of attention after orienting is to modulate sensory processes for perception according to behavioral goals. The attention-related sensory sampling process (attentional sampling, for short) selectively increases the signal-to-noise ratio of the intended sensory information of the oriented target to gain access and form perception (Hillyard et al., 1973; Ungerleider, 2000; Vogel & Luck, 2002). Recently, a rhythmic attention theory (Fiebelkorn & Kastner, 2019) was proposed and hypothesized that the alternating states between attentional sampling and shifting are tethered to neural oscillatory activity. Specifically, the phase of neural oscillations determines different attentional states -- the ‘sampling’ state is assumed in the half period containing peak (termed as the ‘good’ phase), whereas the ‘shifting’ state in another half period containing trough (the ‘poor’ phase). The origin of the neural oscillatory activity that controls the rhythmic attention has been assumed in certain brain regions, such as the pulvinar nucleus of the thalamus (Boshra et al., 2025; Halassa & Kastner, 2017; Kastner et al., 2020). Considering the rhythmic nature of the motor system and the direct link between motor and sensory systems (Assaneo et al., 2019; S. Li et al., 2020), can the motor system induce the modulatory function of attention for shaping perception? Specifically, would the phase of the rhythmic motor activity causally influence attentional sampling?

This study investigates whether and how the rhythmic characteristics of the motor system regulate temporal attention in the auditory domain. We adapted the auditory attentional blink (AB) paradigm to examine the effect of the motor system on attentional process (Shen & Alain, 2010, 2012; Shen & Mondor, 2006). In the AB experiment, participants discriminate a first target (T1) and detect a subsequent probe (T2) in a rapid serial presentation of visual or auditory stimuli. Participants typically display a deteriorated performance on detecting T2 within 200–500 ms after the presentation of T1, regardless of sensory input modality (Arnell, 2006; Dux & Marois, 2009). The AB effect (Broadbent & Broadbent, 1987; Raymond et al., 1992; Vogel & Luck, 2002) is attributed to the transient deficits in the allocation of selective attention (Marti et al., 2015; Sergent et al., 2005; Wyart et al., 2012). We leverage the critical temporal window of AB to directly test the hypothesis that attentional sampling could be driven by phase alignment with rhythmic signals from the motor system.

Across two experiments, we introduced rhythmic manipulations to causally test the relation between motor activity and attentional sampling. In Experiment 1, we manipulated the temporal alignment of the overt movement of rhythmic finger tapping with AB. According to our hypothesis, we predict that the temporal alignment between tapping and T2 presentation would alleviate the auditory AB effect. To investigate whether the phase of the rhythm in motor signals plays a causal role in attentional sampling, in Experiment 2, we applied transcranial alternating current stimulation (tACS) to induce rhythmic neural fluctuations in the target brain areas (Herrmann et al., 2013; Lakatos et al., 2019). Previous studies found that the exogenously induced fluctuations over frontal regions modulate spatial attention (Dugué et al., 2016; Fernández & Carrasco, 2020) and non-spatial attention (Brus et al., 2024). According to the theory of rhythmic attention (Fiebelkorn & Kastner, 2019) in combination with our hypothesis of the causal role of rhythmic motor activity in attentional sampling, we predict that tACS on the motor cortex would modulate the auditory AB effect depending on the phase of simulation. Given the potentially different roles of the motor system in the left and right hemispheres in temporal prediction (Morillon & Baillet, 2017), we predict that dissociable effects of the left and right sensorimotor cortex on alleviating AB would be observed.

## RESULTS

### Rhythmic tapping alleviates the attentional blink

To first examine whether and how action influences attentional processing, in Experiment 1, an auditory AB task was conducted concurrently with finger tapping (Figure 1A). Participants were instructed to identify a target (T1) and then detect a probe (T2) in a rapid serial presentation of auditory stimuli (RSAP, Figure 1A & B). The lag between T1 and T2 was manipulated to generate the attentional blink effect (misses of detecting T2). Crucially, participants tapped their fingers in rhythm along with the preceding tempo of tones, so that the probe was synchronized (Peak, at the same time as a tap) or desynchronized (Trough, between taps) to the tapping rhythm (Figure 1C & D). If action modulates attention in a temporally precise manner, different degrees of synchronization between tapping and probe would change the detection performance differently, compared with a baseline performance without tapping.

**Figure 1.**
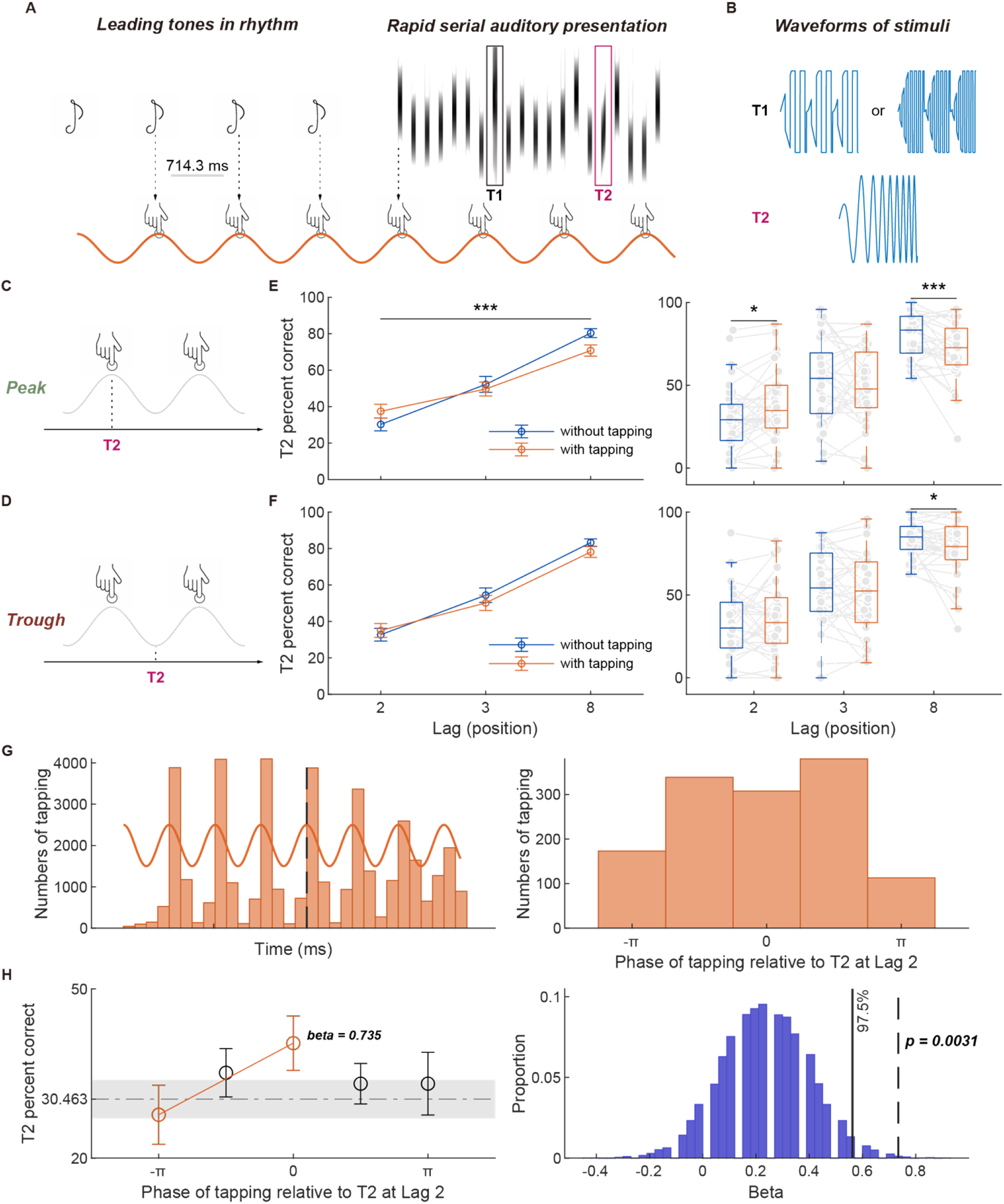
Experimental procedures and behavioral results of rhythmic tapping alleviating the attentional blink. (A) Experimental design. In each trial, four pure tones are presented at 1.4hz, followed by a rapid serial presentation of auditory stimuli (RSAP, SOA of 120 ms). Participants tap their left index finger along with the initial tone rhythm and throughout the trial, at the same time, identify the category of the first target (T1) and detect whether a probe (T2) occurs in the RSAP. The lag between the positions of T1 and T2 is manipulated, and here a lag of 8 is depicted for demonstration. (B) Stimuli examples used as T1 and T2. T1 is composed of eight 5-ms pulses. Two types are yielded by varying the frequency of the pulses. T2 is a glide tone. (C) (C and D) Conditions of temporal alignment between tapping and T2 (C) Peak condition: T2 occurs at the same time as the finger tapping (theoretical timing according to the rhythm of leading tones). (D) Trough condition: T2 occurs in the middle of two taps. (D) (E and F) Behavioral results of T2 performance (detection rate of the probe) as a function of T1-T2 Lag, separately in two alignment conditions as in (C & D). (Error bars represent the s.e.m.) (E) Behavioral performance in the Peak condition. Left, comparison of performance between conditions of different motor involvement across lags. The Motor × Lag interaction was significant, yielding the cross-over of performance in two motor involvement across lags. Right, Results in post-hoc analysis at each lag. Tapping significantly increased the detection rate at Lag 2 but decreased the performance at Lag 8. (F) Behavioral performance in the Tough condition. Left, comparison of performance between conditions of different motor involvement across lags. The Motor × Lag interaction was not significant, yielding no cross-over of performance in two motor involvement across lags. Right, Results in post-hoc analysis at each lag. Tapping only significantly decreased the performance at lag 8. (G) Temporal characteristics of tapping related to auditory stimuli. Left, histogram of timing of taps in the progress of the trial. The dashed line indicates the onset of RSAP. The width of a bin is 178.60 ms. Right, histogram of timing of taps relative to the probe (T2) at Lag 2. The timing is represented as a phase in a cycle relative to the probe, with negative values for taps preceding the probe. Histograms for Lag3 and Lag 8 in Figure S2, and for the timing of taps relative to T1 in Figure S3. The width of one bin is 148.83 ms. (H) Detection of probe as a function of the timing of taps. Left, accuracy on detecting the probe (T2) in the phase bin relative to the probe in Lag 2. The accuracy was highest when the tapping was temporally aligned with the probe (bin of phase 0). A logistic regression revealed a linear increase in the detection rate in the portion of the cycle where taps preceded the probe (bins of negative phase values). The horizontal dashed line represents the baseline level of performance in blocks without tapping; shading represents the s.e.m. Right, the results of a permutation test indicate that the improvement of performance when taps precede the probe is significant. The vertical dashed line represents the empirical value of the correlation in the null distribution.

The detection rate of the probe was subject to a repeated-measures three-way 2 × 3 × 2 ANOVA (Motor involvement × Lag × Phase). The main effects of Lag (F(2, 64) = 122.460, p < 0.001, partial η^2^ = 0.793) and Phase (F(1, 32) = 6.333, p = 0.017, partial η^2^ = 0.165) were significant, but no main effect of Motor (F(1,32) = 0.902, p = 0.304). Critically, a three-way interaction was significant (F(2, 64) = 3.279, p = 0.044, partial η^2^ = 0.093). Following, two-way interaction of Motor × Lag was significant (F(2, 64) = 8.727, p < 0.001, partial η^2^ = 0.214), but not in the other two two-way interactions, Motor × Phase (F(1,32) = 0.156, p = 0.695) or Phase × Lag (F(1,32) = 2.614, p = 0.081). Post-hoc analysis revealed that the significant interaction was because Motor and Lag significantly interacted in the Peak condition (F(2, 64) = 11.156, p < 0.001, partial η^2^ = 0.259, Figure 1E left), but not in the Trough condition (F(2, 64) = 2.872, p = 0.064, partial η^2^ = 0.082, Figure 1F left). These results suggest that motor activity can affect auditory attention, conditioned on the timing between action and probe.

Further analysis of the simple effects of Motor at each Lag in the condition of Peak revealed that the mean accuracy of detection was higher at Lag 2 in the With-tapping condition than that in the Without-tapping condition (t(32) = 2.539, p = 0.016, d = 0.442, Figure 1E right). The effect reversed at Lag 8, where accuracy in the With-tapping condition was lower than that in the Without-tapping condition (Wilcoxon signed-rank test, p = 0.001, Figure 1E right). In the condition of Trough, the performance of T2 detection also decreased at Lag 8 when tapping (Wilcoxon signed-rank test, p = 0.024, Figure 1F right). However, no significant difference was observed at Lag 2 between With and Without tapping conditions (t(32) = 0.739, p = 0.465). Performance at Lag 3 remained unaffected by motor engagement, either in Peak (t(32) = −0.649, p = 0.521, Figure 1E right) or Trough (t(32) = −1.356, p = 0.185, Figure 1F right) condition. These results demonstrate that action can enhance attentional processes when resources are limited, but distract when abundant.

To rule out any general benefits of action, the same repeated-measures three-way ANOVA subject on T1 identification revealed a significant main effect of Motor (F(1, 32) = 11.049, p = 0.002, partial η^2^ = 0.257) and a marginally significant interaction of Motor × Lag (F(2, 64) = 3.111, p = 0.051, partial η^2^ = 0.089).

Post-hoc analysis further shows that the T1 identification decreased significantly in the With-tapping condition (t(32) = 3.324, p < 0.01, d = 0.435, Figure S1), contrasting with the enhancement of T2 detection at Lag 2 when attentional blink occurred. Taken together, these results show a double dissociation of the effects of action on attentional process — activation of the motor system can boost attention when resources are limited, whereas the sensory process is distracted by action when attentional resources are intact.

### Temporal alignment between action and probe enhances sensory processing

The inherent temporal variability of active tasks, such as tapping, renders the manipulation of temporal alignment by design imprecise and hence hinders the validity of the observed effects. Therefore, we directly investigated whether the T2 detection was dependent on the phase of recorded tapping (the actual timing of physical realization). Distribution of the tap timing across all participants was depicted in Figure 1G. The temporal characteristics of tap timing showed a similar pattern related to the preceding tempo of tones, but faded with the progress of the trial, demonstrating that participants followed our experimental design. To model the tapping effect on the T2 detection, we first quantified the actual synchronization between probe and tapping. The relative time difference between the closest tap to the probe was calculated and categorized into five bins, representing different phases of tapping. For example, the third bin means the probe onset was coupled to the closest peak of the tapping rhythm, and the negative value means the tap preceded the probe. The distribution of tap timing relative to the probe at Lag 2 was depicted as Figure 1G (for Lag 3 and Lag 8, see Figure S2). Then the probe performance was modeled as a function of the tapping phase (Figure 1H left).

We conducted a logistic regression to model the behavioral performance as a function of the phase of recorded tapping under conditions of different lags. Results revealed a significant relationship between the tapping phase and T2 detection performance only at lag 2 (permutation test, p = 0.0031, Figure 1H right). However, this enhancement effect did not show at lag 3 and lag 8 (permutation test, p > 0.05, Figure S2). As T1 identification was not manipulated to synchronize with the tapping, only decreased performance was found (Figure S3). These results, after controlling for the temporal variability of tapping, further support that action can enhance sensory sensitivity in a temporally precise manner by boosting attention.

When stimulation over the left sensorimotor (lSM) cortex (left two plots), the T2 detection revealed a U-shape as a function of phase-probe relation. The probe performance (T2) at Lag 2 significantly decreased compared to the Sham condition when the probe was at the start of the falling period (bin 3, paired two-sample t tests, *p < 0.05, left 1). At Lag 7, the probe performance significantly improved compared to the Sham condition when the probe was presented in the same time slot (bin 3, paired two-sample t tests, *p < 0.05, left 2).

When stimulating the right sensorimotor (rSM) cortex (right two plots), the probe performance at Lag 2 significantly improved relative to the Sham condition when the probe was at the start of the rising period (bin 1, Wilcoxon signed-rank test, *p < 0.05, right 1). At lag 7, the probe performance significantly improved compared to the sham condition when the probe was at the start of the falling period (bin 3, Wilcoxon signed-rank test, *p < 0.05, right 2). The dashed line indicates the probe performance at lag 2 or 7 in the sham condition across all participants, and shading depicts the s.e.m. Across all panels, error bars represent the s.e.m.

### Neuromodulation revealed the effects of motor activity timing on attention

Experiment 1 shows that rhythmic tapping can alleviate attentional blink, which hints that the motor system could generate attention-like functions. However, the observed behavioral effects could be a result of actions that are inevitably manifested in a physical form – the tapping provides a physical timing cue and enhances the performance of detecting the probe. To rule out the physical confound and directly test the causal relations between the motor system and attention, we use transcranial alternating current stimulation (tACS), a technique that can periodically modulate neural activities (Brus et al., 2024; Lakatos et al., 2019), in Experiment 2 (Figure 2A). We applied the tACS stimulation within the RSAP paradigm, yielding two phase conditions relative to T2 – a good-phase condition where T2 occurred in the rising period of the tACS stimulation (from phase 0 to peak), and a poor-phase condition where T2 occurred in the falling period of the tACS stimulation (from phase 0 to trough) (Figure 2B). Moreover, the stimulation was separately applied on the sensorimotor cortex over the left or right hemisphere (Figure 2C & D). If the motor system provides attentional-like function from different hemispheres and modulates sensory processing in a specific timing manner, the lateral stimulation in different phases would yield distinct influences on changing the attentional blink.

**Figure 2.**
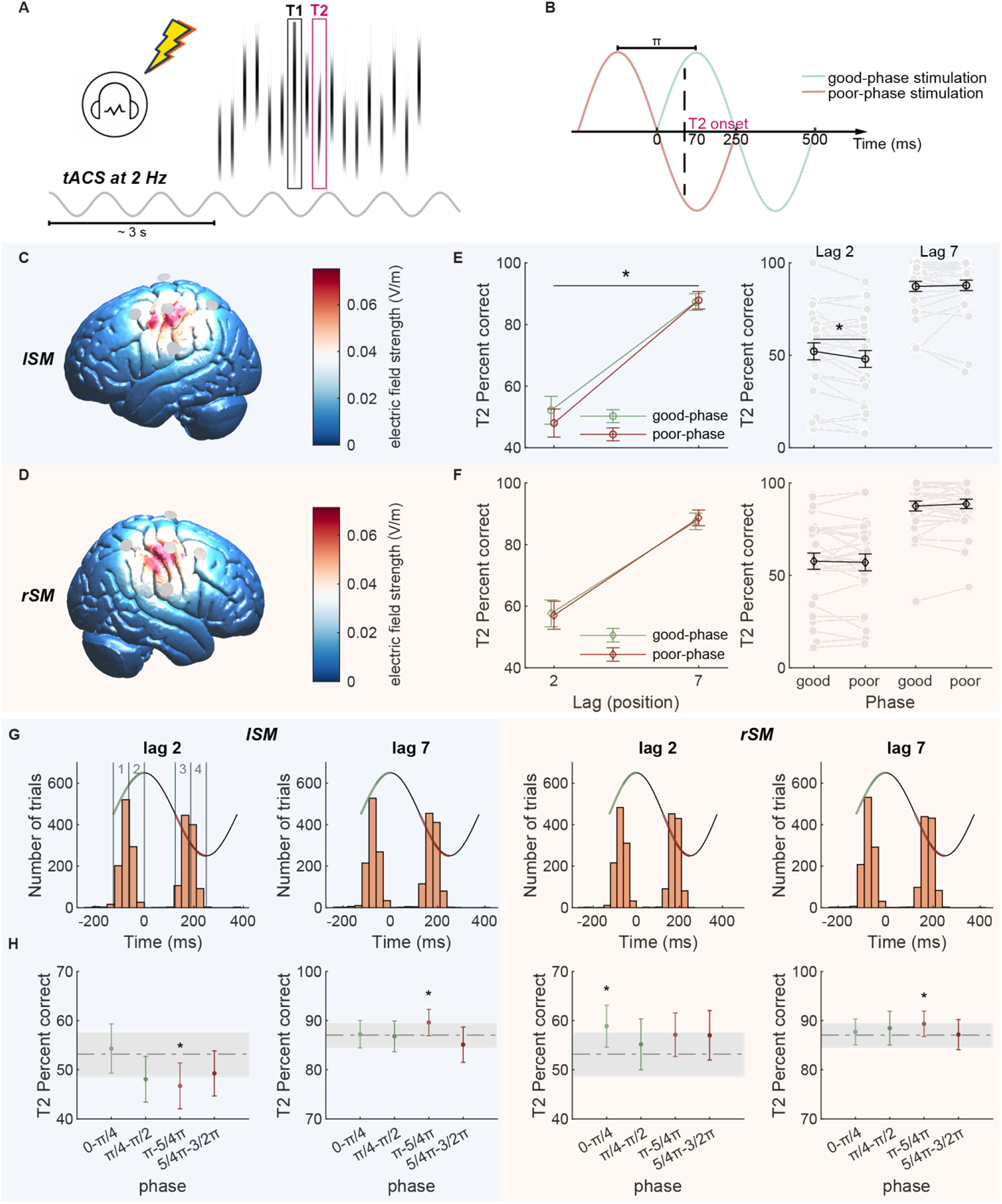
Bilateral sensorimotor cortices differentially modulate attentional blink, revealed by neuromodulation of tACS. (A) Procedures of neuromodulation. Simulation of tACS was applied continuously along the RSAP paradigm. (B) Conditions of stimulation as a function of T2 position. T2 can occur either in the rising period (green line) or in the falling period (red line) of the tACS stimulation, yielding two stimulation conditions of good-phase and poor-phase, respectively (see Methods for details). (C and D) The location of tACS stimulation. The 4×1 array of HD-tACS channels was placed over the left (C, blue panel) or right (D, red panel) sensorimotor cortex. The effective electric fields on the cortical surface were estimated, demonstrating a focal stimulation over sensorimotor cortices. (E) Behavioral performance of T2 detection during stimulation over the left sensorimotor cortex. Left, Detection rate of T2 at Lag 2 and 7, separately in the good-phase (green) and poor-phase (red) stimulation conditions. The phase × lag interaction was significant, suggesting different effects of simulation timing at distinct lags. Right, performance was better in the good-phase than that in the poor-phase stimulation at Lag 2. Individual data are marked by dots connected by lines. (* p < 0.05). (F) Behavioral performance of T2 detection during stimulation over the right sensorimotor cortex. Presentation format is the same as in (E). No significant interaction between phase and lag, nor differences between stimulation phases, was observed. (G) Histograms of the probe onset (T2) relative to the timing of stimulation current. The distribution is presented for Lag 2 and 7, separately for the simulation over the left (lSM) or right (rSM) sensorimotor cortex. Trials were categorized into four bins based on the T2 position relatively to the peak (labeled as 0 ms) of simulation in a cycle, with two bins in the rising phase to the peak (0 to pi/4 and pi/4 to pi/2 as bin1 and 2) and two bins in the falling phase to the trough (pi to 5/4 pi and 5/4pi to 3/2 pi as bin 3 and 4). (H) The modulatory direction of tACS effects revealed by comparisons to the baseline performance in the sham condition (the horizontal dashed line). The phase modulation effect on the T2 detection rate is presented in the four temporal bins according to the four sub-groups in (G).

When applying stimulation on the left sensorimotor cortex, a repeated-measures two-way ANOVA (Lag × Stimulation-phase) revealed a main effect of Lag (F(1, 27) = 140.727, p < 0.001, partial η^2^ = 0.839), but no effect of Stimulation-phase (F(1, 27) = 2.367, p = 0.136). Critically, the interaction of Lag × Stimulation-phase was significant (F(1, 27) = 4.260, p = 0.049, partial η^2^ = 0.136, Figure 2E left). Post hoc analysis revealed that the T2 detection performance was improved only at lag 2 when tACS stimulation was in the good-phase condition (t(27) = 2.252, p = 0.033, d = 0.426, Figure 2E right).

However, when stimulating over the right sensorimotor cortex, only a main effect of Lag was significant (F(1, 27) = 111.269, p < 0.001, partial η^2^ = 0.805, Figure 2F), but no main effect of Stimulation-phase (F(1, 27) = 0.077, p = 0.783) nor interaction of Lag × Stimulation-phase (F(1, 27) = 1.033, p = 0.318). These results suggest that only modulating the left sensorimotor cortex in a specific timing manner induces attentional-like function and changes the attentional blink.

The same analysis was subject to the performance of T1 identification. We only found a main effect of Lag (F(1, 27) = 4.293, p = 0.048, partial η^2^ = 0.137) when stimulating the left sensorimotor cortex. Post hoc analysis shows a worse performance of T1 identification at lag 2 compared to that at lag 7 (t(1) = − 2.072, p = 0.048, d = −0.173, Figure S5). The results indicate a compromised process on T1 as a repulsion relationship between the target and the probe (Marti et al., 2015).

### Dissociable functions of sensorimotor areas on attentional blink

We further specified how the motor system modulates the attentional system. The results of the comparison between two tACS stimulation phases demonstrate the different effects of activating motor systems in changing the attentional blink. But whether the good-phase stimulation enhanced the performance or the poor-phase hindered the performance? Therefore, we further investigated the modulation direction by comparing the performance in different stimulation conditions to the baseline of sham stimulation. Moreover, the actual timing of stimulation may vary from trial to trial, which reduces the power in the investigation of the effects when averaging. We monitored the timing of tACS by concurrent recording from EEG channels, specified the actual timing of stimulation, and grouped into four bins according to the position of T2 in different phases of stimulation (Figure 2G). This precise temporal alignment plus the comparison with the baseline performance, allows us to further specify how the rhythmic motor activity modulates the attentional system.

When stimulation was applied over the left sensorimotor cortex, the performance of T2 detection at lag 2 changed in a U-shape as a function of the phase-probe relation. Specifically, the detection rate at the start of the falling period (pi to 5/4pi of the tACS stimulation, bin3 as in Figure 2G blue panel) was significantly worse than the baseline performance in the sham condition (t(27) = −2.181, p = 0.038, d = − 0.412, Figure 2H left 1). Whereas, at lag 7, the probe performance was reversed into improvement in the same bin (Wilcoxon signed-rank test, p = 0.031, Figure 2H left 2).

When stimulation was applied over the right sensorimotor cortex, the performance of detecting the probe at lag 2 had a trend of elevated accuracy. The boost of performance reached a significant level when the probe appeared in the beginning period of the rising phase (0 to pi/4 of the tACS stimulation, bin 1 as in Figure 2F) (t(27) = 2.426, p = 0.022, d = 0.458, Figure 2H right 1). Whereas, at lag 7, the performance was also improved when the probe appeared in the first period of falling phase (pi to 5/4pi of the tACS stimulation, bin3 as in Figure 2G) (Wilcoxon signed-rank test, p = 0.041, Figure 2H right 2), similarly as observed at lag 7 when stimulation over the left sensorimotor cortex. These results suggest that how the motor system modulates attention co-depends on the timing of the motor signals relative to the sensory processes and the origin of the signals from each hemisphere.

## DISCUSSION

By manipulating the motor engagement that rhythmically covaried with the rapid presentation of stimuli in an attentional blink paradigm, we revealed a causal role of the rhythmic motor activity in the auditory attentional process. The overt movement of rhythmic finger tapping, when temporally aligned with the auditory serial presentation, alleviated the attentional blink effect. Moreover, undulated current stimulation of tACS over the left sensorimotor cortex hindered the attentional process when the probe was presented in the trough of the simulation phase; whereas the detection rate of the probe was facilitated in the peak phase of the stimulation over the right sensorimotor cortex. The consistent behavioral and neural modulation results suggest a causal role of the motor system in regulating the processes of attentional sampling.

Behaviorally, our results show that motor activity can shape attentional processing. The alleviation of the attentional blink by rhythmic tapping, specifically when the probe was synchronized with tapping at Lag 2, indicates that motor signals can rescue attentional processing under resource competition. Prior work has shown that overt rhythmic movements improve detection of temporally predictable auditory events and sharpen auditory temporal selection (Morillon et al., 2014; Morillon & Baillet, 2017; Zalta et al., 2020). The active sensing framework proposes that this occurs through temporal prediction signals – efference copy and corollary discharge – that act in concert with attention to parse rhythmic sensory inputs (Morillon et al., 2015; Schroeder et al., 2010). These studies demonstrate the motor functions in temporally shifting attention. The present study, by situating motor engagement within an attentional blink paradigm, which specifically indexes the failure of selective attention under resource depletion (Dux & Marois, 2009; Sergent et al., 2005; Vogel & Luck, 2002), directly demonstrates that motor signals interact specifically with the attentional sampling process. Together, multi-aspect of evidence supports a complete picture of rhythmic motor activity in orienting and modulatory functions of attention.

Furthermore, the double dissociation of motor effects across lags supports our hypothesis and rules out a general temporal effect. Motor engagement improved T2 detection at Lag 2, but impaired the detection at Lag 8. The T1 identification was also impaired in the movement condition. A general facilitation account would predict uniform benefits regardless of positions. However, the sign of the motor effect reversed across different lags, which suggests that the motor effects on attention depend on whether attentional resources are under competition. Within the serial bottleneck framework (Marti et al., 2015; Sergent et al., 2005; Wyart et al., 2012), the identification of T1 taxes central attentional resources, especially during the blink window, leaving the probe less detectable. Under these conditions, a temporally precise motor signal can act as a priority override – redirecting the gain in attentional sampling toward the temporally-coupled probe. When no such competition exists, such as at Lag 8 or when T1 is under processing, the same motor signal interferes with sensory encoding rather than facilitating. The dissociation between enhancement at Lag 2 and impairment at Lag 8 and T1 suggests that the motor system specifically modulates the attentional sampling mechanism, rather than a general temporal effect.

The behavioral effects observed in Experiment 1 could be caused by physical timing cues that tapping introduced, rather than directly from motor activity. In Experiment 2, we resolved the confounding factor by replacing the overt tapping with direct neural modulation – tACS directly modulating neural oscillations over sensorimotor cortices without any overt movement or external timing cue. The phase-dependent modulation of attentional blink was again observed, which supports that the origin of attention may stem from the sensorimotor neural activity. This causal link is consistent with evidence that tACS-induced oscillatory entrainment modulates attentional performance rhythmically (Brus et al., 2024) and directly demonstrates that the motor system generates attention-like modulation of auditory processing.

Similar motor effects on attentional sampling have also been observed in the visual domain. Subthreshold electrical stimulation on Frontal Eye Field (FEF) causally produced a bottom-up dependent and attention-like modulation effect on visual input and behavioral performance (Ekstrom et al., 2008; Moore & Armstrong, 2003; Moore & Fallah, 2001). Cumulating evidence illuminates the receptive field re-mapping between oculomotor control and spatial attention deployment from behavioral to circuit level (Boshra & Kastner, 2022; Raposo et al., 2023; Rizzolatti et al., 1987; Snyder et al., 1997; Zhou & Desimone, 2011). The motor attentional-like effects in the visual domain primarily depend on the spatial correspondence between the eye movement motor receptive field and the visual receptive field. The current study provides evidence in the auditory domain, further illustrates that the temporal and phasic relations between the rhythmic motor signals and sensory processes mediate the attentional effects across time. Together, the motor system, depending on its temporal and spatial characteristics, can generate attentional-like effects on modulating perception.

We found double dissociated results of a phase-asymmetric effect during the left sensorimotor stimulation. At Lag 2, the probe detection was significantly impaired when the probe appeared at the start of the falling period (approaching trough), while good-phase stimulation did not induce a significant change compared with the sham condition. The phase-specific impairment suggests that the left sensorimotor cortex constrains rather than facilitates attentional sampling during the blink: declining excitability at the critical moment disrupts probe processing. Strikingly, the same phase-probe relationship at Lag 7 reversed into a significant improvement compared with sham. This lag-dissociation effect indicates that the influence of left sensorimotor activity depends on the state of attentional resource, consistent with its role in encoding temporal predictions via delta and beta oscillations directed toward the auditory cortex (Morillon & Baillet, 2017). This resource-dependent, phase-specific pattern was, however, specific to the left hemisphere, pointing to dissociable contributions of bilateral sensorimotor cortices in attentional sampling.

Critically, the hemispheric asymmetry between left and right sensorimotor effects provides an important functional dissociation. Previous findings demonstrate lateralized attentional functions in lesion studies (visuospatial hemineglect, see (Cohen et al., 1994; Corbetta & Shulman, 2011)) and functional neuroimaging studies (Corbetta & Shulman, 2002; J. T. Coull & Nobre, 1998). However, we found phase-specific and asymmetric effects stemming from bilateral motor areas for modulating auditory attention. The left sensorimotor cortex produced a phase-dependent detrimental effect at Lag 2. The right sensorimotor cortex, by contrast, produced only facilitatory effects at Lag 2, suggesting a more general excitatory contribution to sensory processing under competition rather than precise temporal gating. Critically, both hemispheres showed performance improvement at Lag 7 when stimulation was at the trough, presumably when the sensorimotor cortex was deactivated. These results showed a consistent pattern with the behavioral finding that motor activity is disruptive when attentional resources are intact. The converging behavioral and neurophysiological results across both lags suggest that the motor influence on attention is resource-dependent and bidirectional, mediated by left and right sensorimotor cortices through distinct but complementary mechanisms. However, whether bilateral cortices serves in a interhemispheric competition way (Szczepanski & Kastner, 2013), simultaneous bilateral neuromodulation with concurrent EEG will be needed to map the interhemispheric dynamics underlying this asymmetry. Moreover, Henry & Obleser (2012) showed that the lag between sensory events and intrinsic brain oscillations is individually variable, suggesting that individualized, EEG-guided stimulation protocols would sharpen the precision of these effects further.

Converging behavioral and neurophysiological results reveal that the motor system exerts a strong influence on attentional processes. However, the observed pattern also presents a challenge to existing theories. The active sensing theory (Schroeder et al., 2010) predicts that motor-derived temporal predictions consistently facilitate auditory processing. However, our data show motor activity is detrimental when attentional resources are intact (Lag 8, T1). Similarly, the rhythmic theory of attention (Fiebelkorn & Kastner, 2019) predicts that activating the motor system competes with already-depleted attentional resources during the blink. However, our data show motor engagement specifically rescues probe detection under the challenging condition. Neither framework could fully account for the observed bidirectional, resource-dependent motor effects.

We propose an asymmetrical gain modulation derived from the interaction between motor signals and attentional resource allocation (attentional state). This asymmetrical gain modulation is based on two inherent properties of motor internal signals: (1) predictive function and (2) bottom-up dependent modulation. The first is a general predictive function: the motor system generates temporally precise signals via efference copy and corollary discharge, which is well established across predictive coding theory (Parr & Friston, 2019), internal forward models (S. Li et al., 2020; Tian & Poeppel, 2010; Wolpert et al., 1995), and causal inference frameworks (Rao, 2024), and have been shown to stabilize sensory representations (Moore et al., 1998; Sommer & Wurtz, 2006) and support learning (Tourville & Guenther, 2011). The second property is a bottom-up dependent directional bias: previous work shows that FEF microstimulation in non-human primates modulates early visual areas (V1-4) only when the bottom-up sensory process is activated, where gain modulation is reversed between early and higher-order visual areas and the strongest modulation effect is in early visual areas (Ekstrom et al., 2008; Moore & Armstrong, 2003).

Applying the asymmetrical gain modulation in the current study, when attentional resources are scarce during attentional blink (internal state), the motor prediction carried no identity content (i.e., no acoustic content was predicted), but was temporally precise and coupled to the probe (external stimulus). The temporal prediction generates a timing prediction error upon probe arrival. This unsuppressed prediction error may be further boosted by the second property, which is the directional bias, so that the enhanced prediction error may reset the attention network by tuning up the gain of attentional sampling toward the probe. When no such competition exists, the same motor signals interfere with ongoing sensory encoding. That is, motor signals serve as a bottom-up dependent regulator of attentional sampling, which implies that the top-down modulation from the motor system may function in parallel with bottom-up attention. Thus, in the interaction between the internal state and external world, motor signals may be another source, like working memory (Nobre & Gresch, 2025) that actively support the orienting and sampling functions of attention.

In sum, combining behavioral manipulation and sensorimotor neuromodulation in an auditory attentional blink paradigm, our findings reveal the motor system as an asymmetrical, resource-dependent regulator of attentional sampling, with the left sensorimotor cortex gating access in a phase-dependent manner and the right providing a more general competitive boost. These results inspire a new direction for understanding how the motor system shapes cognitive functions in a rhythmic manner.

## RESOURCE AVAILABILITY

### Lead contact

Further information and requests for resources and reagents should be directed to and will be fulfilled by the lead contact, Xing Tian.

### Materials availability

The current study has not generated any new material.

### Data and code availability

The code associated with this paper is available at: https://osf.io/hnz96/

## ACKNOWLEDGMENTS

We thank Yunying Shu, Xizi Wang and Shiya Cheng for their assistance in data collection, and Lizzie Gao, Xinjing Li, Zhili Han and Yuchunzi Wu for their help in experimental design and analysis. This study was supported by the National Natural Science Foundation of China (32271101) to X.T. and (32471136) to Y.K., and Program of Introducing Talents of Discipline to Universities Base B16018, and NYU Shanghai Boost Fund to X.T.

## AUTHOR CONTRIBUTIONS

Conceptualization, Z.X., Y.L. and X.T.; methodology, Z.X., J.S., Y.L., W.Z., Z.W., Y.K. and X.T.; Investigation, Z.X., Y.L. and W.Z.; writing—original draft, Z.X.; writing—review & editing, Z.X., Y.K. and X.T.; funding acquisition, Z.W., Y.K. and X.T.; supervision, Y.K. and X.T.

## DECLARATION OF GENERATIVE AI AND AI-ASSISTED TECHNOLOGIES

During the preparation of this work, the authors used Claude to improve the readability and language of the manuscript. After using this tool or service, the authors reviewed and edited the content as needed and take full responsibility for the content of the publication.

## METHODS

### EXPERIMENTAL MODEL AND STUDY PARTICIPANT DETAILS

Forty-five (age range: 18–26 years; 22 females) and forty (age range: 18–25 years; 20 females) volunteers participated in Exp. 1 and 2, respectively. All participants had normal audition and vision, and reported no history of neurological or psychiatric disorders. A short survey before the experiments indicated that none of the participants were professional musicians. Twelve and ten participants were excluded from analysis in Exp. 1 and 2, respectively, because they had poor performance or no attentional blink effects (see QUANTIFICATION and STATISTIC ANALYSIS). Another two participants were excluded in Exp. 2, because of instrumental errors of tACS during experimentation. Hence, the total participants were 33 and 28 in Exp. 1 and 2, respectively. Informed consent was obtained from all participants before the experiments. All experiments were approved by the local ethics committees at East China Normal University and NYU Shanghai.

### METHOD DETAILS

#### Stimuli

In Exp. 1, auditory stimuli were sampled at 44.1 kHz using Adobe Audition 2023. All these stimuli were adapted from the auditory attentional blink (AB) task (Shen & Mondor, 2006). This task includes three categories of auditory stimuli – target (T1), probe (T2) and distractors. T1 was composed of eight 5-ms pulses (Figure 1B shows the 15-ms examples of T1 and T2). The pulses were either 529 or 1270 Hz, yielding two tokens for T1. T2 was a tone glide and its frequency changed continuously from 636 to 1006 Hz. Twenty-one pure tones were used as distractors. The frequencies of these pure tones were logarithmically sampled from 529 to 1330 Hz. All auditory stimuli were 40 ms in duration, including a 2-ms linear onset/offset amplitude ramp. Different from the original AB task, a pure tone at 918 Hz was repetitively presented 4 times to construct a leading rhythm before the AB task. The duration of this pure tone was 71.4 ms. The pure tone was ramping up/down in 2 ms. In Exp. 2, auditory stimuli in the AB task were changed to 30 ms duration to obtain a more stable attentional blink effect.

#### tACS settings

We used a current stimulator (NeuStim, Neuracle) with a manual stimulation protocol controlled by TXCS software (Neuracle Inc.). The 4×1 HD-tACS setting was followed the montage used by Spooner & Wilson (2023), in addition that we positioned the electrodes (outer radius = 1.2 cm, inner radius = 0.6 cm) on both sides of M1 (i.e., central electrode: C3, surrounding electrodes: C1, C5, FC3, CP3; central electrode: C4, surrounding electrodes: C2, C6, FC4, CP4). To proximately approach the rhythmic tapping task as Experiment 1, tACS was applied at a frequency of 2 Hz (period of 0.5 s). To create two approximately equivalent circuits and minimize skin sensation, the skin was prepared with conductive gel while keeping the net impedance below 5 kΩ. Electric field simulation on a standard head model (Montreal Neurological Institute [MNI] template) revealed a relatively focal distribution of field intensity over the target regions using SimNIBS toolbox (SimNIBS v4.5.0). Figure 2C and D depict the theoretical current intensity of this electrode montage with a stimulation intensity of 0.5 mA (Thielscher et al., 2015).

To minimize the potential confounding effect from fluctuating somatosensory inputs, we determined the amplitude of each participant during a pre-testing session. In this session, the amplitude was set individually by adjusting the peak amplitude of the current, separately for each montage in a step of 0.1 mA, starting from 0.5 mA to the point where participants reported feeling uncertain about the presence of tACS under every electrode (0.562±0.117 mA average across participants, as in Li et al. (2023)). At the beginning of each block, the current was ramped up over the first 10 s. For the blocks applying sham stimulation, we ramped up the current to its maximum over 10 s and turned it off as a baseline condition. The tACS was applied continuously during the stimulation blocks and monitored by EEG recordings simultaneously.

#### EEG Data acquisition

EEG was recorded using a 32-channel active electrode system (Brain Vision actiCHamp; Brain Products), with a sampling rate of 1000 Hz in an electromagnetically shielded and soundproof room. The electrode arrangement was according to the international 10–20 system. The impedance of each electrode was kept below 10 kΩ. The data were referenced online to the electrode of Cz. The EEG data were acquired with Brain Vision PyCoder software (http://www.brainvision.com/pycorder.html). Because of the massive stimulation artifact introduced from the tACS stimulation, the contaminated EEG data were primarily used to monitor the actual tACS current relative to triggers induced by auditory stimuli (Figure S6).

#### Procedures of Experiment 1 Behavioral task

The behavioral paradigm is depicted in Figure 1A. The experiment was conducted in a dimly illuminated room. Instructions were displayed on a mid-grey background on a screen with a spatial resolution of 1920 by 1080 pixels and a vertical refresh rate of 60 Hz. Auditory stimuli were presented binaurally at a comfortable hearing level via headphones or via plastic air tubes connected to foam earplugs (Sennheiser HD 280 linear in Experiment 1 and ER-3C Insert Earphones; Etymotic Research in Experiment 2), using the Psychtoolbox-3 and additional custom scripts written for MATLAB (The Mathworks). Timing of participants’ tapping was recorded by a trackpad (perixx pp504 linear) and participants were instructed to tap lightly to minimize somatosensory feedback.

Each trial consisted of two stages – a leading rhythm stage and a rapid serial auditory presentation (RSAP) stage. The leading rhythm comprises four pure tones presented in the stimulus-onset-asynchrony (SOA) of approximately 714.3 ms. The RSAP stage was composed of twenty auditory stimuli (SOAs: 120 ms). The target (T1) was at one of two positions (5 or 8) in the RSAP to reduce temporal expectation, which can reduce the AB effect (Shen & Alain, 2012). The probe (T2) was presented at a probability rate of 75% at one of three possible positions relative to T1 (2, 3, 8 with respect to the occurrence of the target (T1), thereafter referred to as lag 2, 3 and 8; Factor 1, *Lag*). All other stimuli in the RSAP were distractor sounds.

Participants were instructed to perform a dual-task experiment in the RSAP stage. The first task was to discriminate one of two targets at T1 (see stimuli). The second task was to detect the probe. At the end of each trial, the response screen for the first question (‘Please type the sort of target [A] or [L]?’) appeared around 500 ms after the auditory stimuli. The second question (‘Was there a Tone Glide? yes: 1; no: 0?’) appeared following the response to the first question. Participants were required to report by pressing the corresponding buttons. They were asked to respond as accurately as possible.

Moreover, the probe occurred in phase or anti-phase relative to the theoretical tapping time determined by the preceding leading rhythm (Factor 2, *Phase*). For example, when the probe was lag 2 or 8 following the target at the 5th position of the RSAP, the onset of T2 was temporally coupled to the peak of the leading rhythm. However, when the onset of T2 was lag 2 or 8 following the target at the 8th position of the RSAP, the probe was temporally coupled to the trough of the leading rhythm. In addition, when the probe was lag 3 to the target, the probe was 120 ms away from the peak and trough, which was the control condition to evaluate the pure effect of motor over the effect of phase.

Finger tapping was the third factor that could accompany the presentation of an auditory stimuli sequence (Factor 3, *Motor involvement*). In the With-tapping condition, participants were required to follow the leading rhythm with their left index finger on a noiseless trackpad from the beginning of the sequence to establish a tapping rhythm and continued to tap in the rhythm during the following RSAP sequence. An encouragement (‘Please try to follow the Leading rhythm∼’) would be presented when tapping records were fewer than three within one trial. In the Without-tapping condition, participants were asked to stay still and minimize any overt involvement.

Together, our experimental protocol resulted in a 3 × 2 × 2 factorial design with the factors of Lag (2, 3, 8), Phase (peak versus trough) and Motor involvement (with versus without tapping). The experiment was divided into two sessions of two lab visits in one day (at least a 2-hour interval). Each session consisted of trials in the With or Without tapping conditions, separately. The order of With-tapping and Without-tapping sessions was counter-balanced between participants. In each session, subjects performed 6 blocks (32 trials each). The presentation order of trials in a block was pseudo-random. In total, 24 trials per condition were acquired with 384 trials in total.

Participants practiced before the main experiment. Participants first listened to the auditory stimuli of the target (T1) and the probe (T2) to facilitate memorizing and discriminating; meanwhile, the response keys were associated with the corresponding auditory stimuli (‘A’ to the 529 Hz target, ‘L’ to the 1270 Hz target and ‘1’ to the probe). Before the With-tapping session, a tapping practice block was conducted to familiarize participants with the tapping rhythm. Participants were required to tap their left index finger following a rhythm that was the same as the leading rhythm in the main experiment. A pure tone was played back via the headphones as the feedback of tapping. Participants were encouraged to keep up with the rhythm and maintain a steady rhythm. The length of the tapping practice trial was 20 s. At least three practice trials were conducted, with more practice trials upon request by participants.

Before the main experiment, two more practice blocks were conducted to familiarize participants with the procedures of the experiment. The procedures were identical to the main experiment of RSAP except that 1) only 12 trials per block were included and 2) feedback on the discrimination of T1 was provided. The practice was conducted before the with-tapping and without-tapping sessions.

#### Procedures of Experiment 2 tACS task

The experimental procedures of Exp. 2 are depicted in Figure 2A. The procedures are similar to those in Exp. 1, with a couple of exceptions. First, because the performance of probe detection was similar at Lag 2 and 3 in Exp. 1, to save time, only Lag 2 and 7 were included in Exp. 2. Second, the stage of leading rhythm was excluded to avoid the effects of auditory entrainment. Instead, tACS stimulation was started 3 s before the RSAP. Two stimulation conditions, *good-phase stimulation* and *poor-phase stimulation*, were included. According to the results of Exp. 1, manual action can facilitate the detection of the probe only when tapping preceded the probe in a range of 0 to 120 ms (Figure S4). Therefore, we set 70 ms as the critical window between auditory stimuli and motor activation. During the *good-phase stimulation*, the probe preceded the peak of electric stimulation by 70.884±30.213 ms (measured by simultaneous EEG recordings, Figure S6). During the *poor-phase stimulation*, the phase of the current was shifted by 180º. The stimulation conditions were pseudo-random on a trial-to-trial basis.

The stimulation was applied at two locations, either at the left or right sensorimotor cortices (Figure 2C and D). We applied the 4×1 HD-tACS centered at the location of C3 according to the international 10-20 system (above the left cerebral hemisphere with surrounding electrodes: C1, C5, FC3, CP3) and C4 (above the right cerebral hemisphere with surrounding electrodes: C2, C6, FC4, CP4). Moreover, a sham condition was included as a baseline condition, where the current ramped up over the first 10 s to its maximum and then ceased before each sham block. The order of stimulation sites and sham was randomized across blocks, except for the first 5 participants, where the order of stimulation montages was counter-balanced. The protocol resulted in a 2 × 2 factorial design with factors of *lag* (lag 2 and 7), *stimulation-phase* (good-phase stimulation vs. poor-phase stimulation), separately in two stimulation sites (left and right sensorimotor cortices).

The experiment was divided into two lab visits, separated by at least three days. Participants completed 13 and 12 blocks in each visit, respectively (24 trials each block). In total, 600 trials were conducted with 40 trials per condition. A similar practice to that in Exp. 1 was provided in Exp. 2. In the first visit, a behavioral pre-test was conducted, including a training and a practice session. The training session was the same as Exp. 1 for familiarizing the two targets and the probe. The practice session included three blocks with each of 24 trials that were identical to the main experiment. Feedback was provided in the first practice block. In the second visit, participants performed one training session to refresh their memory with the auditory targets and one main experiment.

### QUANTIFICATION AND STATISTICAL ANALYSIS

#### Scores of behavioral performance and the magnitude of the attentional blink effect

The accuracy (percent correct) was obtained in the discrimination task of T1 and the detection task of T2. In Exp. 1, participants whose discrimination performance on T1 deviated by more than 3 standard deviations from the group and whose performance of T2 at lag 8 < 50% (suggesting nearly random choice selection and therefore indicating poor engagement) were excluded from data analyses, resulting in the exclusion of five participants. In Exp. 2, Participants were excluded from further analysis if their performance of T1 in the sham condition deviated more than 3 standard deviations from the group mean, resulting in the exclusion of one participant.

In Exp. 1, the magnitude of the attentional blink (AB) effect was calculated using the following formula as in Willems et al. (2013):

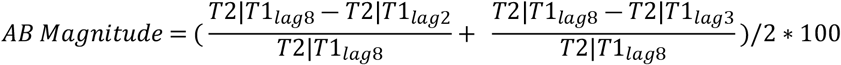

T2|T1_lag_ is the mean accuracy of probe (T2) at a specific lag, given that target (T1) was correctly reported. The AB magnitude is quantified as the averaged difference between behavioral performances of probe (T2) outside (lag 8) and inside (lag 2 and 3) the critical AB window. Specifically, we calculated the difference between the accuracy at Lag 8 and the average accuracy of Lags 2 and 3, divided by the accuracy at Lag 8. This formula compares the probe (T2) performance in the critical time window (200-500 ms) relative to the one out of the time window of attentional blink. To avoid erroneously categorizing participants who had overall low T2 accuracy as non-blinkers, we used T2|T1_lag8_ instead of T1_mean_.

In Exp. 2, because we have two levels of lags for T2 position relatively from T1 (lag 2 and lag 7), AB magnitude was calculated as follows:

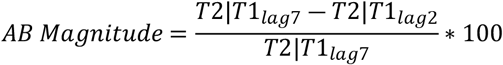

Participants with AB magnitude < 10% were excluded from further analysis because they had no auditory attentional blink effects, resulting in the exclusion of seven and nine participants in Experiments 1 and 2.

### Behavioral analysis

All data analyses were conducted in MATLAB R2021a (MathWorks) using custom scripts. Statistical tests were computed using JASP (JASP Team (2024)). To examine the effects of motor activity on the auditory attentional process, we conducted a repeated-measures three-way 2 × 3 × 2 ANOVA (Motor involvement × Lag × Phase), with the conditional probability of T2 detection rate given a correct T1 discrimination as the dependent variable. Follow-up paired t-tests were conducted to assess the motor effect on auditory AB at different lags. Moreover, the T1 performance was subject to the same three-way ANOVA and post-hoc t-tests. When normality was violated, the Wilcoxon signed-rank test was used.

To evaluate the relationship between the recorded taps and the detection performance of T2, we used logistic regressions to model the probe performance as a function of the tapping phase. First, the relative time difference between the recorded tapping and the probe onset was calculated in each trial. The closest tapping to the probe onset was used in the following analysis. Secondly, according to the leading rhythm (as the rhythm of theoretical tapping was the same as the leading rhythm), we divided the tapping into 5 phase bins (bin = 148.833 ms) centered around the probe onset. Each tapping event was assigned to a phase bin based on the relative timing of the tapping to the probe. Finally, we performed a logistic regression analysis using a Generalized linear model (GLM). We modeled the T2 performance as a function of the tapping phase category (the first and third bin). The model was fitted to the pooled data across 31 participants for each Lag condition (Lag 2, 3, and 8). Two participants were excluded from this analysis because they lacked recorded tapping in the first or the third bin. We extracted the beta coefficients from the models to quantify the weights of the modulation effect of different phases on T2 detection. Statistical significance was tested by a permutation test with 10,000 iterations. In each iteration, we shuffled the correspondence between the tapping phase labels and the behavioral performance within each subject to preserve individual variability while changing the phase-behavior relationship. The shuffled data were fitted to the same logistic regression model to generate a null distribution of beta coefficients. P-value was calculated by determining the position of the empirical beta value in the null distribution. Modeling and parameter estimates were generated using the MATLAB fitglm function and custom scripts.

#### Analysis of tACS effects

To causally examine the effects of the motor system on the auditory attentional process, we conducted two repeated-measures two-way 2 × 2 ANOVAs (Lag × Phase) on the T2 performance, separately for the tACS stimulation over the left and right sensorimotor cortex. Follow-up paired t-tests were conducted between different phase conditions in different lags. T1 performance was also subject to the same analyses. When normality was violated, the Wilcoxon signed-rank test was used.

#### Phase-dependent effect analysis

To precisely determine the phase of tACS, we simultaneously recorded EEG signals. Firstly, we selected Cz as the index electrode because it is strongly affected by the tACS and phase equilibrium between the two electric fields centered at C3 and C4. Then we extracted an epoch from −250 to 250 ms (half cycle of 2 Hz tACS), relative to the probe onset for each trial. Thereafter, we labelled and indexed the phase of tACS stimulation by identifying the local maximum (peak) and minimum (trough) of the energy fluctuation within an epoch. The actual phase of the tACS was obtained by calculating the temporal difference between the onset of the probe and the nearest peak or trough of the stimulation current. The validity of the stimulation was confirmed via onset slope analysis, leading to the removal of 1.2% of trials (about 5.32 trials per participant) as artifacts.

To examine the phase effect of sensorimotor cortex stimulation on the auditory AB, we further divided each trial into four sub-groups. Specifically, we used a 65 ms (i.e., a phase width of 1/4 π in the 2 Hz tACS) time window to bin each trial according to the temporal distance between the onset of the probe and the peak or trough of the current stimulation, yielding two bins preceding the peak and two bins preceding the trough. The T2 performance was calculated as a function of each bin. The phase effect of stimulation was statistically tested by comparing it to those in the sham condition using paired t-tests. When normality was violated, the Wilcoxon signed-rank test was used.

